# Antibody Complementarity-Determining Region Sequence Design using AlphaFold2 and Binding Affinity Prediction Model

**DOI:** 10.1101/2023.06.02.543382

**Authors:** Takafumi Ueki, Masahito Ohue

## Abstract

Affinity maturation in the immune response is limited in terms of affinity gain, and natural antibodies often do not have the binding affinity required for therapeutic applications. Therefore, improving the binding affinity of antibodies is essential for developing antibody-based therapeutics. Designing antibodies using experimental methods is expensive in terms of cost and time owing to the large range of complementarity-determining regions to be explored. Recently, computational methods have been developed as low-cost and fast means of designing and redesigning antibodies. This study evaluated the design performance of AlphaFold2 and the binder hallucination, which can predict protein 3D structures with high accuracy even without experimental antibodies, by redesigning antibody sequences to improve the binding affinity of existing antigen-antibody complexes. Therefore, antibody sequences with higher affinity can be designed for antigen-antibody complexes not included in the training data of AlphaFold2, indicating that the proposed method may be effective as an antibody design method.

## I. Introduction

Affinity maturation of human and non-human natural antibodies in the immune response is limited by the limited increase in their binding capacity [1]–[4]. Therefore, in many cases, they do not have the binding ability necessary for therapeutic purposes. Improving the binding affinity is an important step in developing antibody drugs.

The structure of an antibody comprises two regions: a variable region and a constant region. In the variable region, the loop structure called the complementarity-determining region (CDR) is important for recognizing antigens. Although non-CDRs are composed of relatively similar amino acid sequences, the CDR has diverse sequences and structures. Therefore, it is possible to generate antibodies with high specificity for antigens because they can assume various shapes.

Generating antibodies with high specificity for antigens involves arranging amino acid sequences in the variable region. The CDR comprises six regions, L1 to L3 and H1 to H3, and each amino acid sequence length is approximately 4–30 residues. When selecting sequences with high affinity and specificity using experimental methods, the number of sequence combinations to be examined is enormous, resulting in exorbitantly high costs in terms of money and time.

In recent years, with advances in computational technology, various computational methods have been developed in protein engineering, attracting much attention because they reduce experimental costs. In antibody design, language model-based methods in deep learning can efficiently generate antibody sequences that bind well to antigens compared to experimental methods in terms of experimental costs. However, language model-based methods are based on sequence-based methods. Therefore, they are unsuitable for designing mutations in sequences with high variabilities, such as CDR or sequences without experimental data [5]–[7].

In protein engineering, remarkable progress has been achieved in the field of conformation prediction, particularly with the development of AlphaFold2 [8], which can predict protein conformations with high accuracy. Recently, a technique called “hallucination” that applies the ability to accurately predict protein structures to protein design has gained attention [9].

In this study, we used the AfDesign protein design method [10], which uses AlphaFold2 for structural prediction in hallucination, to redesign the amino acid sequences in the CDR as the design site for existing antibody-antigen complexes. We aimed to take advantage of AlphaFold2’s ability to predict protein structures with high accuracy, even when experimental structures were unavailable.

## II. Methods

### A. Binder hallucination

In this study, we focused on the binder hallucination function of AfDesign and applied it to the antibody sequence design problem. A schematic of the methodology used in this study is shown in Fig. 1. Binder hallucination designs protein sequences using the following mechanism [11]:

**Fig. 1:**
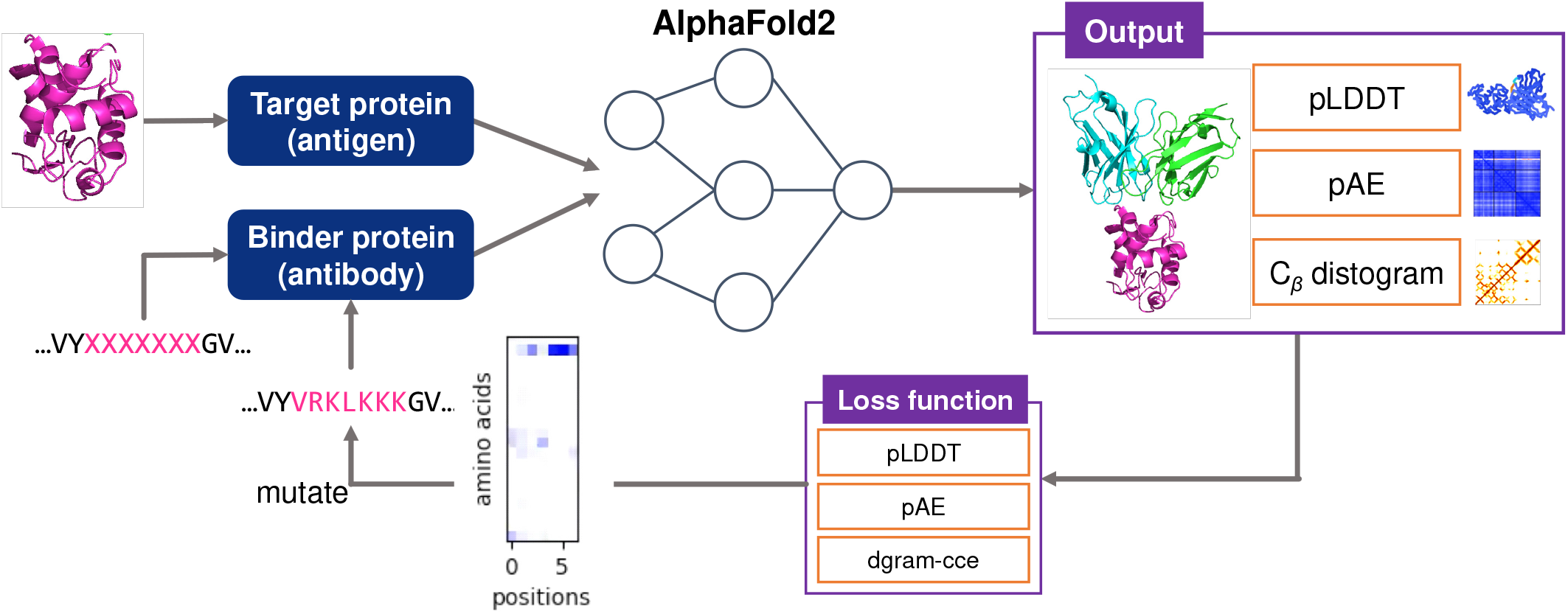
Schematic of binder hallucination.

1. Specify the type of target protein and the sequence length of the binder protein.
2. The amino acid sequence of the binder side is randomly generated at the specified length.
3. AlphaFold2 predicts the 3D structure of the target protein-binder protein complex and outputs pLDDT and pAE, which represent the reliability of the prediction and the C_*β*_-distogram, a distance matrix between C_*β*_ atoms, respectively.
4. The amino acid sequence is updated to minimize the loss using dgram-cce, pLDDT, and pAE as loss functions (dgram-cce is a categorical cross-entropy of the distogram).

Binder hallucination is a protein design approach that simultaneously “generates a three-dimensional structure” and “searches for a sequence” by iteratively repeating the processes described in steps 3 and 4, aiming to design a protein that is expected to bind to a target protein.

In this study, a stepwise optimization method (`design_3stage()`) was provided by AfDesign to directly optimize the one hot encoded array when dealing with complex topologies, such as three-dimensional structures. This method involves optimizing the output of the neural network before passing it through the softmax activation function (logits), followed by optimizing the output of the softmax activation function (soft), and finally optimizing the array encoded in the one-hot representation (hard). This approach allows optimization to be performed for complex topologies [12] and was employed in this study.

In the binder hallucination function of AfDesign, existing proteins can be partially redesigned by preparing the amino acid sequence of the binder protein and specifying the site to be redesigned in Step 1. Therefore, in this study, we applied binder hallucination by replacing the target with an antigen and the binder with an antibody and targeted the CDR amino acid sequences of the antibody as sites for the redesign. We used AfDesign to design sequences with higher binding affinities than those of the original sequences.

### B. Output value of AlphaFold2

#### 1) pLDDT

The pLDDT metric is an adaptation of the lDDT index introduced by Mariani *et al*. [13]. The lDDT metric is commonly used to assess the accuracy of predicted protein structures in terms of their ability to replicate corresponding reference structures. Specifically, it involves the calculation of all interatomic distances between a set of atoms belonging to a target residue in the reference structure and a set of atoms within a predetermined distance threshold (*R*_0_) that do not belong to the same residue. The percentage of atoms within the threshold that are also present in the predicted structure is computed to quantify the extent to which the predicted structure preserves the local geometry of the reference structure. The pLDDT index builds on this metric by incorporating additional factors that account for uncertainties and errors inherent in the prediction process, resulting in a more robust and reliable measure of structural accuracy.

AlphaFold2 calculates lDDT-C_*α*_, which targets only the C_*α*_ carbons in each residue, and trains it to output pLDDT, which is the predicted value of lDDT, even in cases where the correct structure is unavailable. It outputs the reliability of the prediction for each residue within a range of 0 to 100. The regions with low pLDDT values often correspond to intrinsically disordered regions that do not have specific structures [14]. In AfDesign, pLDDT is expressed as a value between 0 and 1.

#### 2) pAE

pAE is a matrix representing the positional error between the C_*α*_ atoms of residue *j* and residue *i*, which is calculated by aligning the predicted structure by AlphaFold2 using the backbone structure of residue *i* with the experimentally determined structure. When predicting structures for which no ground-truth structure is available, regions where the relative error between residues *i* and *j* is large indicate lower confidence in the prediction [15].

#### 3) C_β_ distogram

In AfDesign, the distance matrix (distogram) representing the distance between C_*β*_ and C_*β*_ residues predicted by AlphaFold2 can be used. Using dgram-cce as a loss function allows the design of predictions that have higher confidence in forming complexes [16].

### C. Valuation index

In this study, the change in binding free energy during the antigen-antibody binding process, ΔΔ*G*, was used to indicate the strength of the antigen-antibody interaction. However, because it is difficult to obtain the value of ΔΔ*G* experimentally, we used the value predicted by the calculation in this study. In this study, we used the DDG Predictor [17], [18], a deep learning-based ΔΔ*G* prediction tool.

## III. Experiments

### A. Experiment 1: Determination of Loss Weights

In the binder hallucination function of AfDesign, the dgram-cce, pLDDT, and pAE described in Section II-B can be used as loss functions, and AfDesign allows the user to set the weight of each loss. In this experiment, we varied the relative weights of dgram-cce, pLDDT, and pAE losses to redesign the antibody sequences under different conditions and investigated the appropriate weights of these losses.

#### 1) Protein Design Tools

AfDesign, a protein design tool available at the GitHub repository [10], was used in this study. Software version 1.0.8 was used in this study.

#### 2) Various parameters

The following parameters were used in the experiment: the number of iterations represents the number of iterations of logits, soft, and hard in the `design_3stage()`. In AfDesign, the number of times that AlphaFold2 iteratively improves the structure through cycling is specified as num recycle.

- Design object: 1VFB
- Number of repetitions: logits-soft-hard=50-50-5
- num recycles = 0
- Learning rate: 0.01
- Optimization Algorithm: Adam [19]

Ten samples were generated for each of the six regions of the CDR, L1–3, and H1–3 under six conditions (A—F). For each of the 360 samples generated, the values of ΔΔ*G* were outputted using the DDG Predictor. CDRs were determined according to Chothia’s definition [20]. The coordinates of atoms other than the target atom were fixed, and structure prediction using AlphaFold2 was performed only for the target amino acid.

Case A dgram-cce=1.0

Case B dgram-cce=1.0, pLDDT=0.2, pAE=0.2

Case C dgram-cce=1.0, pLDDT=0.4, pAE=0.4

Case D dgram-cce=1.0, pLDDT=0.6, pAE=0.6

Case E dgram-cce=1.0, pLDDT=0.8, pAE=0.8

Case F dgram-cce=1.0, pLDDT=1.0, pAE=1.0

#### 3) Valuation index

Using ΔΔ*G* predicted by the DDG Predictor, the average ΔΔ*G* and IMP (IMProved Percentage), the percentage of sequences generated with a higher binding capacity than the original amino acid sequence was used as the valuation index.

### B. Experiment 2: Application of AfDesign to existing antigen-antibody complexes

Based on the results of Experiment 1, we determined the appropriate weights for the losses pLDDT, pAE, and dgram-cce and applied AfDesign to the CDRs of the 12 complexes manually selected from the Protein Data Bank (PDB) in Table I. For each CDR, we generated 10 samples by redesigning with AfDesign and randomly mutating the amino acids. A total of 720 samples were generated using these two methods, and the predicted ΔΔ*G* values were obtained by inputting the mutant and original PDB files into the DDG Predictor for each sample.

**TABLE I:** Protein Data Bank (PDB) IDs of 12 antigen-antibody complexes (complexes in bold were not used for AlphaFold2 training [8])

|  |
| --- |
| 1QNZ, 1VFB, 1JHL, 2DQC, 2YSS, 2UZI,<br>4XVJ, 5NGV, 6J5F, **7CWO**, **7RR0**, **7WP1** |

In Experiment 2, the following parameters were used:

- Number of repetitions: logits-soft-hard = 50-50-5
- num recycles = 0
- Learning rate: 0.01
- Optimization Algorithm: Adam [19]
- Types of loss functions and their respective weights: dgram-cce=1.0, pLDDT=1.0, pAE=1.0

As in Experiment 2, Using ΔΔ*G* predicted by the DDG Predictor, the average ΔΔ*G* and IMP, the percentage of sequences generated with a higher binding capacity than the original amino acid sequence were used as valuation indexes.

## IV. Results and Discussion

### A. Experiment 1: Determination of Loss Weights

Fig. 2 shows the distribution of ΔΔ*G* values for 60 samples when designing the six regions of CDR-L1–3 and CDR-H1–3, each with 1VFB, as design targets under various conditions. According to Fig 2, there is no difference in the distribution among the different conditions; no advantage of ΔΔ*G* by condition was found.

**Fig. 2:**
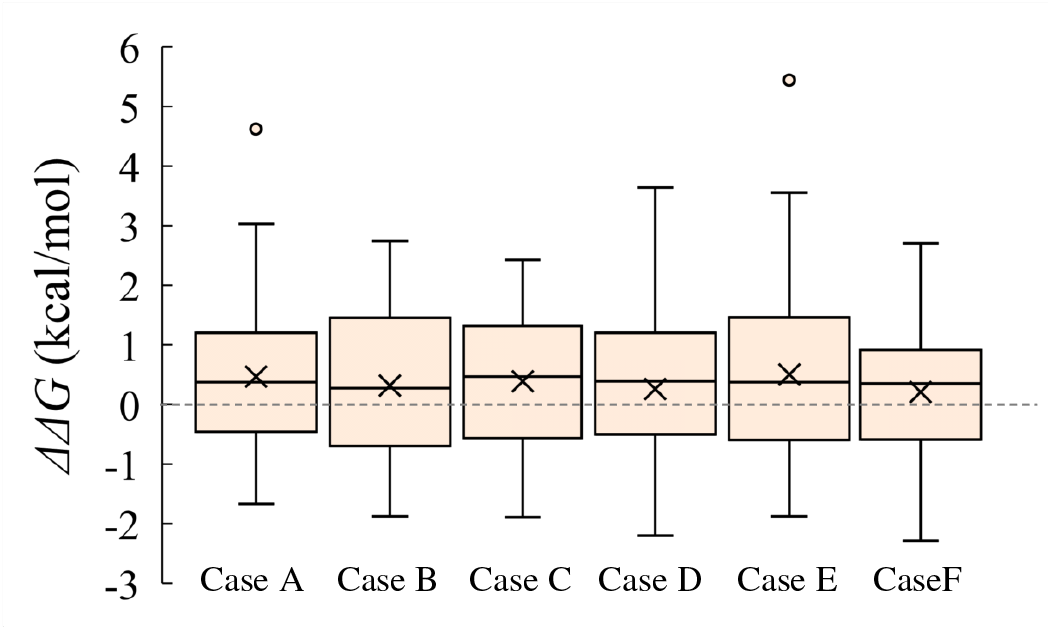
Distribution of ΔΔ*G* for each condition

Fig. 3 is a graph that shows the transition of the three parameters, dgram-cce, pLDDT, and pAE, during the design process under various conditions when designing CDR-H1 as the design target. Looking at the transition graph for Case A, even though only dgram-cce was considered a loss, pLDDT and pAE also showed correlated changes in their values. In addition, even when the proportions of pLDDT and pAE in the loss function increased in Cases B and C, the changes in the transitions of the two parameters were very small.

**Fig. 3:**
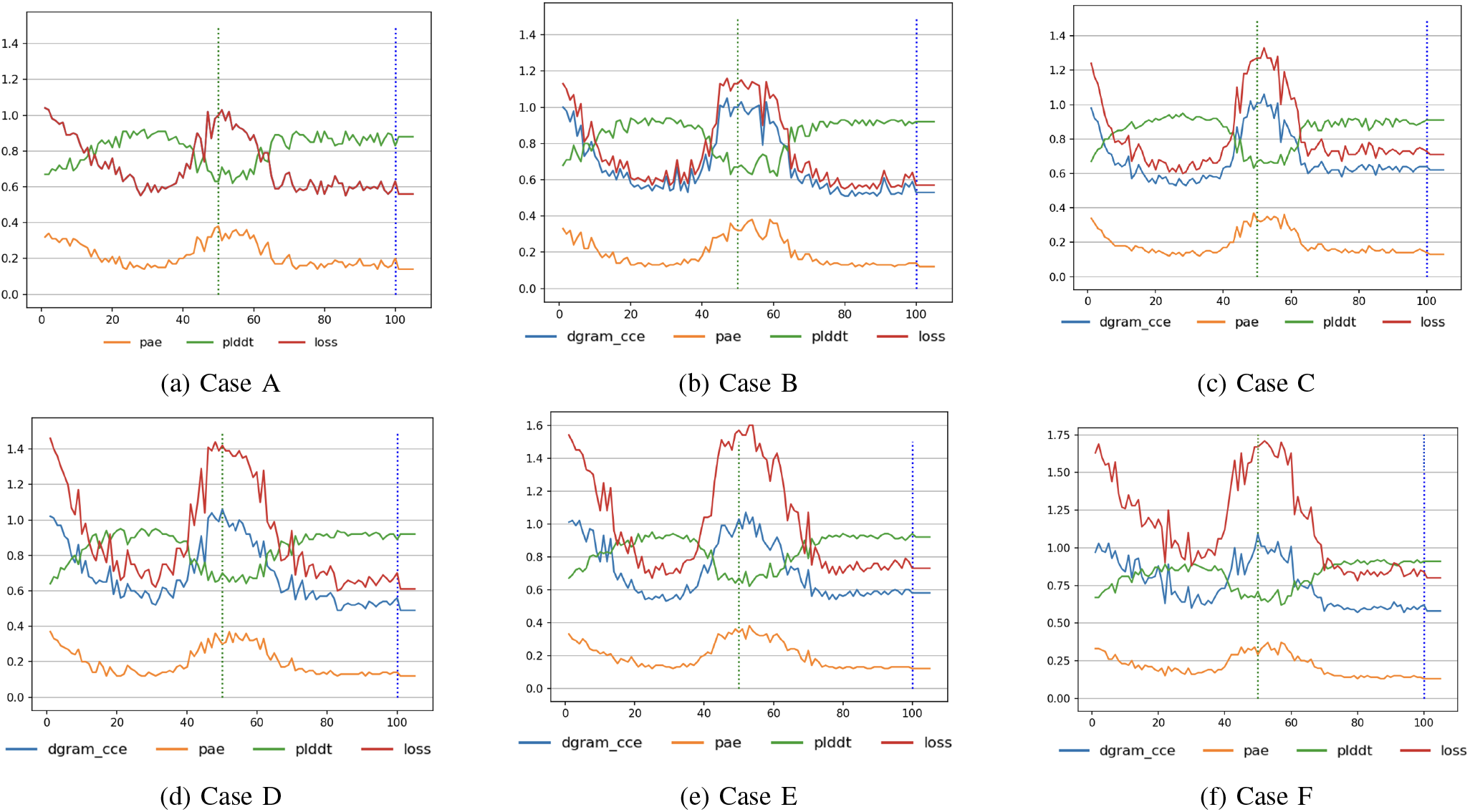
1VFB CDR-H1 as the design target and the evolution of the values of the three parameters when designed for each condition.

Table II lists the average ΔΔ*G* values for the ten samples generated under each condition when each CDR was the design target, as well as the average ΔΔ*G* and IMP values for each condition. Although no clear advantage in binding affinity was found depending on the conditions, we conducted Experiment 2 using Case F, which showed the best average ΔΔ*G* and IMP values.

**TABLE II:** Average ΔΔ*G* and IMP values (PDB 1VFB) for each condition. Letters in bold indicate the best value in each CDR.

|  | Case A | Case B | Case C | Case D | Case E | Case F |
| --- | --- | --- | --- | --- | --- | --- |
| CDR-L1 | -0.797 | **-0.816** | -0.420 | -0.549 | -0.666 | -0.699 |
| CDR-L2 | 1.096 | 1.046 | 0.935 | 1.063 | 0.926 | **0.829** |
| CDR-L3 | -0.561 | -1.020 | -1.385 | **-1.480** | -0.795 | -0.833 |
| CDR-H1 | -0.255 | -0.306 | -0.084 | -0.114 | -0.376 | **-0.414** |
| CDR-H2 | 1.547 | 1.279 | 1.555 | 1.231 | 1.430 | **0.945** |
| CDR-H3 | 1.787 | 1.702 | 1.713 | 1.409 | 2.491 | **1.404** |
| avg $\Delta\Delta G$ | 0.46 | 0.31 | 0.38 | 0.26 | 0.50 | **0.20** |
| IMP (%) | 45.0 | **46.7** | 36.7 | 43.3 | 45.0 | **46.7** |

### B. Experiment 2: Application to existing antigen-antibody complexes

Fig. 4(a) shows the comparison between designs obtained using AfDesign and those obtained from 20 random mutations, in which 20 amino acids were randomly mutated with equal probability within the same design range. In particular, when CDR-H1 and CDR-H3 were targeted for design, more sequences with higher binding affinity than the original sequence were generated compared to random mutations. However, when CDR-L2 and CDR-H2 were targeted, the design performance was inferior to random mutations, and the results differed depending on the CDR.

**Fig. 4:**
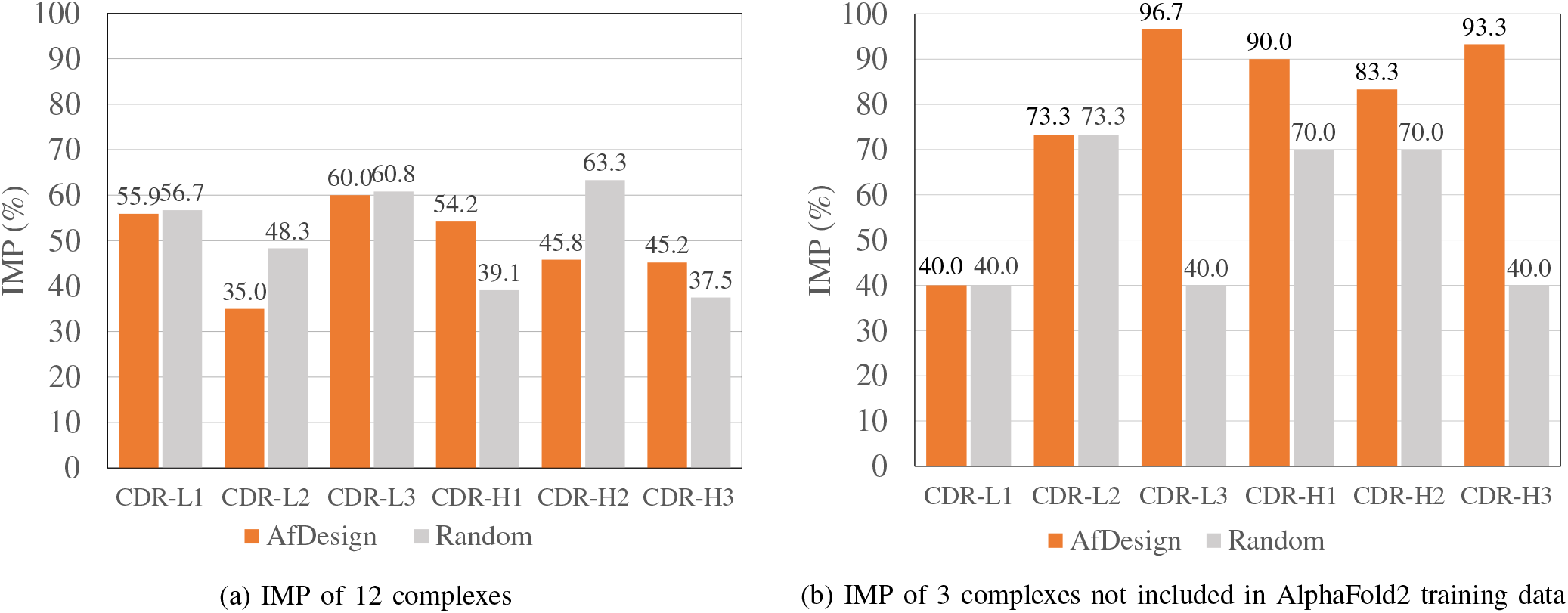
IMP of sequence design results for each of AfDesign and Random mutation

Fig. 4(b) shows the results for three of the 12 complexes used in Experiment 3 that were not used for AlphaFold2 training. It was confirmed that in all regions, a higher percentage of sequences with higher binding affinity than the original sequence was generated compared to random mutations or at least the same level. Thus, it is possible to create mutations that improve binding affinity, even without training data.

Figs. 5(a) and 5(b) show the superimposed 3D structures of the actual sequences generated using AfDesign with CDR-L1 and CDR-H3 as the design targets and the original structures for PDB ID 7RR0. The version of AfDesign used in this study utilized the proteins registered in PDB in 2019 for AlphaFold’s training; therefore, the 7RR0 complex registered in PDB in 2021 was not used for training. However, when designing regions other than CDR-H3, as shown in Fig. 5(a), there was a tendency to generate sequences with structures similar to the original loop structure, even for complexes not used in AlphaFold2 training. The results of this study show that CDR-H3 tends to produce a structure that is slightly different from the original loop structure, as shown in Fig. 5(b). This is because it is difficult to predict the loop structure of CDR-H3 because of the diversity in the sequence and structure of CDR-H3 compared with those of other regions of the CDR.

**Fig. 5:**
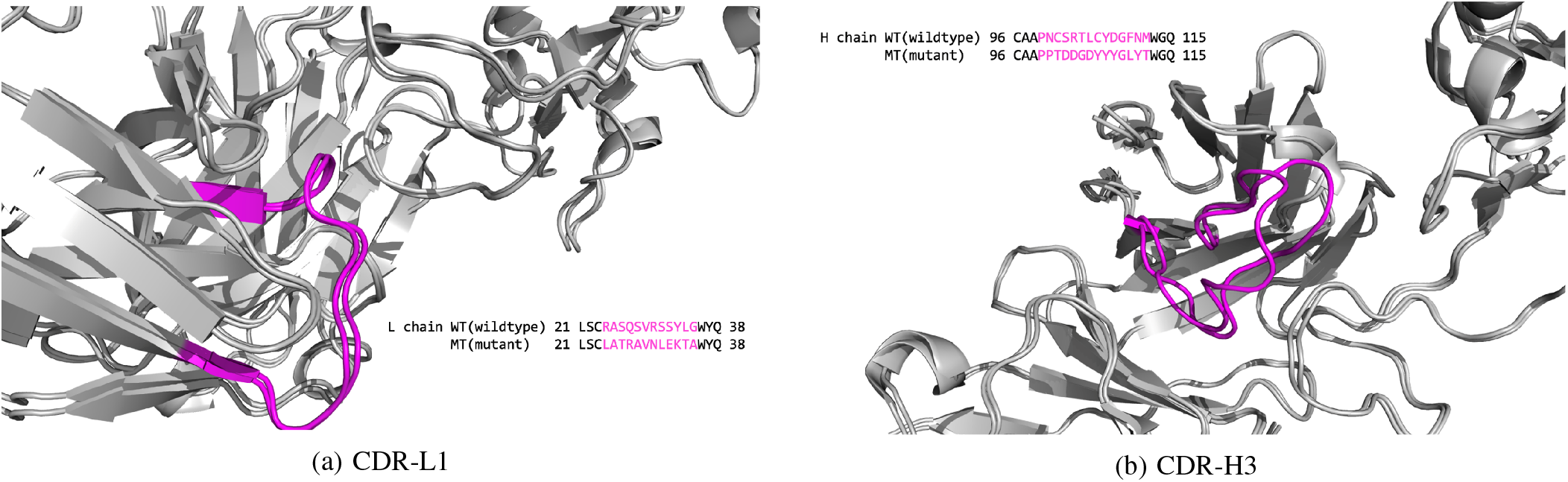
Sequence redesign of PDB ID 7RR0 structure by AfDesign

Therefore, we speculate that the difference in CDR-H3 loop structure prediction accuracy by AlphaFold2 may be due to the generated loop structures being slightly offset from the original loop structures.

## V. Conclusion

In this study, we designed antibody sequences using AlphaFold2, which can predict molecular 3D structures with high accuracy, even without training data. In related studies, there were issues with the design of sequences without experimental data or with highly variable sequences, such as CDRs. However, the results of Experiment 2 in this study confirm that AfDesign can generate sequences that significantly improve binding affinity compared to randomly mutated sequences, particularly in cases without experimental data. This suggests that AfDesign is highly effective for designing sequences with improved binding affinities, even in cases where little or no experimental data are available.

In addition, using dgram-cce, pLDDT, and pAE as loss functions in AfDesign generated sequences that tended to resemble the original loop structure. This was observed even for complexes that were not used in the training of AlphaFold2. Thus, AfDesign can be highly effective in generating CDR loop structures that bind well to unknown antigens; however, further investigation is necessary.

In this study, machine learning prediction tools were used for evaluation. Therefore, it is necessary to verify whether the designed sequences bind to the targets through biochemical experiments.

## Acknowledgment

This work was partially supported by JST FOREST (JP-MJFR216J). We thank Takatsugu Kosugi for the useful discussions.

## References

[1] L. A. Rabia, A. A. Desai, H. S. Jhajj et al., “Understanding and overcoming trade-offs between antibody affinity, specificity, stability and solubility,” Biochem. Eng. J., vol. 137, pp. 365–374, 2018.

[2] M. L. Chiu, D. R. Goulet, A. Teplyakov et al., “Antibody structure and function: the basis for engineering therapeutics,” Antibodies, vol. 8, no. 4, p. 55, 2019.

[3] J. Foote and H. N. Eisen, “Kinetic and affinity limits on antibodies produced during immune responses.” Proc. Natl. Acad. Sci., vol. 92, no. 5, pp. 1254–1256, 1995.

[4] F. D. Batista and M. S. Neuberger, “Affinity dependence of the B cell response to antigen: a threshold, a ceiling, and the importance of off-rate,” Immunity, vol. 8, no. 6, pp. 751–759, 1998.

[5] E. C. Alley, G. Khimulya, S. Biswas et al., “Unified rational protein engineering with sequence-based deep representation learning,” Nat. Methods, vol. 16, no. 12, pp. 1315–1322, 2019.

[6] J.-E. Shin, A. J. Riesselman, A. W. Kollasch et al., “Protein design and variant prediction using autoregressive generative models,” Nat. Commun., vol. 12(1), no. 2403, pp. 1–11, 2021.

[7] J. Linder, N. Bogard, A. B. Rosenberg et al., “A generative neural network for maximizing fitness and diversity of synthetic DNA and protein sequences,” Cell Syst., vol. 11, no. 1, pp. 49–62, 2020.

[8] J. Jumper, R. Evans, A. Pritzel et al., “Highly accurate protein structure prediction with AlphaFold,” Nature, vol. 596, no. 7873, pp. 583–589, 2021.

[9] I. Anishchenko, S. J. Pellock, T. M. Chidyausiku et al., “De novo protein design by deep network hallucination,” Nature, vol. 600, no. 7889, pp. 547–552, 2021.

[10] sokrypton, “AfDesign,” 2023, https://github.com/sokrypton/ColabDesign/tree/main/af.

[11] T. Kosugi and M. Ohue, “Solubility-aware protein binding peptide design using AlphaFold,” Biomedicines, vol. 10, no. 7, 2022.

[12] sokrypton, “af/readme.md,” 2023, https://github.com/sokrypton/ColabDesign/blob/main/af/README.md.

[13] V. Mariani, M. Biasini, A. Barbato et al., “lDDT: a local superposition-free score for comparing protein structures and models using distance difference tests,” Bioinformatics, vol. 29, no. 21, pp. 2722–2728, 2013.

[14] K. Tunyasuvunakool, J. Adler, Z. Wu et al., “Highly accurate protein structure prediction for the human proteome,” Nature, vol. 596, no. 7873, pp. 590–596, 2021.

[15] Y. Moriwaki, “The trajectories of protein structure prediction before AlphaFold2 and the future,” JSBi Bioinform. Rev., vol. 3, no. 2, pp. 47–60, 2022.

[16] J. P. Roney and S. Ovchinnikov, “State-of-the-art estimation of protein model accuracy using AlphaFold,” Phys. Rev. Lett., vol. 129(23), no. 238101, 2022.

[17] S. Shan, S. Luo, Z. Yang et al., “Deep learning guided optimization of human antibody against SARS-CoV-2 variants with broad neutralization,” Proc. Natl. Acad. Sci., vol. 119, no. 11, p. e2122954119, 2022.

[18] HeliXonProtein, “DDG Predictor,” 2023, https://github.com/HeliXonProtein/binding-ddg-predictor.

[19] D. P. Kingma and J. Ba, “Adam: A method for stochastic optimization,” arXiv preprint arXiv:1412.6980, 2014.

[20] C. Chothia and A. M. Lesk, “Canonical structures for the hypervariable regions of immunoglobulins,” J. Mol. Biol., vol. 196, no. 4, pp. 901–917, 1987.

